# An updated evolutionary and structural study of TBK1 reveals highly conserved motifs as potential pharmacological targets in neurodegenerative diseases

**DOI:** 10.1101/2022.09.16.508274

**Authors:** Louis Papageorgiou, Eleni Mangana, Eleni Papakonstantinou, Io Diakou, Katerina Pierouli, Konstantina Dragoumani, Flora Bacopoulou, George P Chrousos, Themis P Exarchos, Panagiotis Vlamos, Elias Eliopoulos, Dimitrios Vlachakis

## Abstract

TANK binding kinase 1 protein (TBK1) is a kinase that belongs to the IκB (IKK) family. TBK1, also known as T2K, FTDALS4, NAK, IIAE8 and NF-κB, is responsible for the phosphorylation of the amino acid residues Serine and Threonine. This enzyme is involved in various key biological processes, including interferon activation and production, homeostasis, cell growth, autophagy, insulin production and the regulation of TNF-α, IFN-β and IL-6. Mutations in the *TBK1* gene alter the protein’s normal function and may lead to an array of pathological conditions, including disorders of the Central Nervous System. The present study sought to elucidate the role of the TBK1 protein in Amyotrophic Lateral Sclerosis (ALS), a human neurodegenerative disorder. A broad evolutionary and phylogenetic analysis of TBK1 was performed across numerous organisms to distinguish conserved regions important for the protein’s function. Subsequently, mutations and SNPs were explored and their potential effect on the enzyme’s function was investigated. These analytical steps, in combination with the study of the secondary, tertiary, and quaternary structure of TBK1, enabled the identification of conserved motifs, which can function as novel pharmacological targets and inform therapeutic strategies for Amyotrophic Lateral Sclerosis.

## Introduction

Amyotrophic lateral sclerosis (ALS) is one of the rarest neurological diseases that affects adults, manifesting as the progressive loss of both lower and upper motor neurons (LMNs and UMNs) (1). On a neurological level, the nerve cells that control the muscles, such as the spinal cord (lateral) and the brain, are destroyed and sclerosis is precipitated. As a result, the brain is unable to activate and control the movement of muscles (2, 3). Respiratory failure is the main cause of fatal outcomes in ALS patients and may occur within three to five years from the onset of symptoms (3).

According to epidemiological data of 2019, the prevalence is 4.1-8.4 / 100,000 people, while worldwide the incidence is 0.6-3.8 / 100,000 people, with Europe having the highest incidence and amounting to 2.1-3.8 / 100,000 (3). ALS can be divided into two categories, familial ALS (fALS) and sporadic ALS (sALS). The sporadic type of ALS constitutes 90 – 95 percent of total cases and is unrelated to family history of the disease (4). The majority of the risk for sporadic ALS is genetically determined and while a variety of environmental factors are suspected of involvement, they remain unproven conclusively (5). Familial ALS accounts for the remaining 5 –10 percent of total cases and is linked to a specific gene (6). The general age period for the appearance of symptoms is 50 years and 60 years for familial and sporadic ALS respectively (7). ALS is less prevalent in women compared to men, with a frequency of 1:1.5, however in familial ALS, the frequency of prevalence appears equal between males and females (8). Half of ALS patients do not survive the two-year mark after the onset of disease and more than 10 percent of patients will live for ten or more years (2).

In the case of familial ALS, mutations have been found to play a crucial role in its emergence. At least twenty genes harbor mutations and most incidents of fALS and frontotemporal dementia (FTD) are caused by mutations in the *C9orf72* gene (9). fALS-related mutations have also been detected in other genes, such as TARDBP, FUS, UBQLN2 and SOD1 (10). Aside from genetic ones, environmental factors such as metals, solvents and radiation are under investigation as potential ALS risk factors (11). Neurotoxin L-BMAA, contained in fish, mussels, and oysters, has been detected in the brains of patients who died from ALS (12). Work exposure to dangerous environmental toxins such as pesticides and lead may enhance the risk of developing ALS, as evidenced in the case of military veterans (13).The onset of ALS may be triggered by an array of dysfunctions, such as mitochondrial dysfunction, dysregulation of glutamate metabolism, oxidative stress, protein aggregates and accumulation of neurofilaments (2). Early symptoms of ALS onset include - but are not limited to - muscle weakness affecting an arm leg, the neck or the diaphragm, muscle cramps, muscle twitches in the arm, leg, shoulder or tongue, slurred and nasal speech, spasticity and difficulty chewing and swallowing (14).

To this day, there is no available cure for ALS. In order to avoid complications and limit the effect of symptoms on the patient’s everyday life, drugs such as edaravone and riluzole are available (4). Edaravone can slow the progression of ALS (15) and riluzole has been proven to reduce the glutamate levels resulting in decrease of motor neuron injury but bears no effect on the existing damage (16).

The *TBK1* gene, located in 12q14.2, encodes the enzyme TANK binding kinase 1 (TBK1), which is also known as NAK, NF-Kappa-B-Activating Kinase, T2K, FTDALS4 and IIAE8 (17). TBK1 is a member of the non-canonical IkB kinases (IKK) and is responsible for the phosphorylation of nuclear factor kB (NFkB) (18). More specicifically, TBK1 adds phosphates to the oxygen of amino acids Threonine and Serine. Generally, TBK1 participates in many cellular functions, such as apoptosis, cell duplication and growth and autophagy (19). Furthermore, TBK1 can activate interferon regulatory factors 3 and 7 (IRF-3, IRF-7), thereby participating in the induction of IFN-I production and antiviral immunity (20). Changes in *TBK1* expression or loss of TBK1 normal function can contribute to the emergence of pathological conditions such as cancer and a variety of neurodegenerative disorders (21–23).

TBK1 is an 82 kDa protein with a length of 729 amino acids (24). TBK1 is composed of three principal domains: residues 9-301 constitute the kinase domain (KD), residues 305-383 constitute the ubiquitin-like domain (ULD) and residues 407 – 729 constitute the domain with elongated helices (ED) (25). ED encompasses two coiled-coin domains, CCD1 and CCD2, at residues 407-657 and 658-713 respectively, which primarily mediate homodimer formation between TBK1 and IKKi (25). Furthermore, the C-terminus of the CCD2 domain has an adapter capture pattern that facilitates TBK1 interaction with TANK adapters (26). Residues 499-529 belong to a leucine zipper (LZ) inside the CCD1, while a helix-loop-helix domain (HLH) is formed by residues 591 – 632 (**Figure 1**) (26). The KD and ULD domains form the N-terminus of TBK1, while the C-terminus is formed by CCD1 and CCD2 (25). In order for TKB1 to be activated, lysine residues 30 and 401 are required to undergo poly-ubiquitination and a serine residue at position 172 is subsequently required to undergo phosphorylation (19).

**Figure 1:**
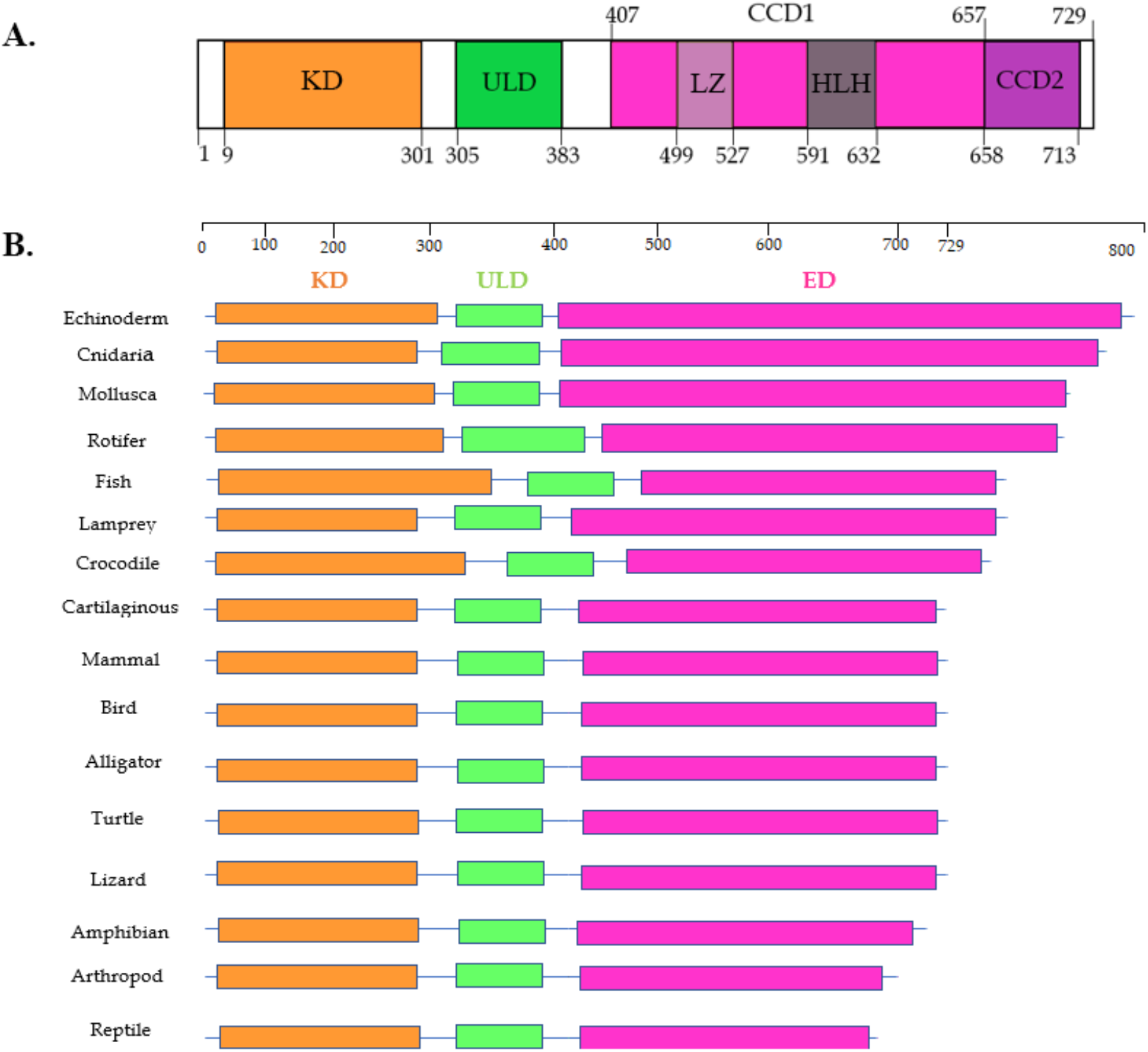
**(A)** Domains of human protein TBK1 and its length 729 aa. **(B)** Multiple sequence alignment (MSA) of protein TBK1 expressed in organisms and the appearance of conserved and nonconserved domains. The orange domain is the kinase domain (KD), the green domain is the ULD, and the pink domain is the domain with the elongated helices (ED). Large conservation is observed at the KD and ULD, with small exceptions, whereas the ED domain exhibits low conservation between the different species.

The *TBK1* gene is involved in many biological pathways and as a result, mutations occurring in the gene can have implications in a variety of pathological conditions. Typical examples include Frontotemporal Dementia (FTD), childhood Herpes Simplex Encephalitis (HSE), Amyotrophic Lateral Sclerosis (ALS), Normal Tension Glaucoma (NTG) and Primary Open Angle Glaucoma (POAG) (19, 27–29). The effect of mutations on the protein’s function heavily depends on the domain where the mutation occurs. Mutations in the kinase domain can potentially damage the site of TBK1 phosphorylation, in the ULD domain may affect the binding site, in the CCD1 domain may hinder TBK1 activation and mutations in the CCD2 domain may affect adapter-TBK1 connection (26). Generally, the kinase domain and CCD1 are sites with high probability of harboring mutations (19). Mutations in ULD and CCD1 are more likely to cause ALS, while mutations in CCD2 are more likely to cause ALS-FTD or FTD (26, 30–32). In conclusion, TBK1’s implication in normal and diseased physiological states renders the enzyme a particularly attractive subject of research. In the context of ALS, the study of multiple aspects of TBK1, including its evolution, genetic variation and structure, can elucidate the enzyme’s role in the disorder and guide the development of novel therapeutic approaches.

## Methods

### Dataset collection and filtering

In order to accumulate the protein sequencies for analysis, the NCBI database (ncbi.nlm.nih.gov) was queried using keywords and synonym names for TBK1, such as protein NAK, T2K, NF-Kappa-B-Activating Kinase, FTDALS4 and IIAE8. The protein sequences were selected from all TBK1-expressing organisms and the results were stored in FASTA format for each taxonomic group separately, creating separate datasets. The datasets were subsequently filtered with the use of MATLAB Bioinformatic Toolbox in order to eliminate potential noise in the form of data unrelated to the protein of interest (33). Protein sequences that responded to the query but did not include TBK1 were eliminated from the primary dataset, by using related keywords and regular expressions techniques in the header information and the FASTA format (34). Irrelevant proteins, putative, low-quality proteins, predicted, isoforms, partial sequences and various transcription factors (cofactor, coregulator, coactivator) were filtered out of the datasets with appropriate MATLAB scripts. Lastly, protein sequences in each species dataset that were found to share more than 95% protein identity within each dataset were removed (35, 36).

### Multiple sequence alignment, conserved motifs and phylogenetic analysis

Multiple sequence alignment (MSA) was executed using the MATLAB Bioinformatics Toolbox. The progressive multiple alignment was carried out with the help of a guide tree, as previously described (37). Pairwise distances among sequences were estimated based on the pairwise alignment with the “Gonnet” method, followed by calculation of the differences between each pair of sequences (38). The Neighbor-Joining method was implemented to estimate the guide tree, assuming equal variance and independence of evolutionary distance estimates. The aligned TBK1 sequences were subsequently studied carefully in Jalview for the identification of conserved regions (39). The commentary section of Jalview, which presents the amino-acid conservation using logos and histograms, was further observed to uncover novel motifs (40). Lastly, phylogenetic analysis was performed with MATLAB Bioinformatics Toolbox, utilizing the Unweighted Pair-Group Method (UPGMA) (41). The constructed phylogenetic tree was visualized using the MEGA radiation option, and the final clusters were separated by different colors (42).

### Single nucleotide polymorphisms, variants, and mutation analysis

Mutations and single nucleotide polymorphisms (SNPs) can have a strong impact on gene function and by extension, on protein function (43). Therefore, their study can yield interesting results regarding genetic variation in the context of health and disease. Information regarding human TBK1 genetic variation was collected through the use of publicly available databases and scientific literature. Typical examples of these sources include the Protein Data Bank (PDB), Uniprot, GWAS Catalog and Pubmed Database (44–46). A collection of currently known naturally occurring TBK1 mutations and SNPS were assembled following a survey of the existing scientific literature. The selected mutations exhibited detrimental effects on the activation and the phosphorylation of TBK1. The resulting dataset includes naturally occurring TBK1 mutations in the following domains: kinase domain, ULD domain, CCD1 and CCD2 domain. Based on the generated list, mutations were identified and marked on the consensus sequence in the study’s multiple sequence alignment (40).

### Structural analysis

The structural and functional analysis of human TBK1 was performed by collecting and studying crystal structures through the Protein Data Bank (PDB) (33). Limiting the search to structures of human TBK1, two crystals were used, PDB ID: 5W5V and PDB ID: 6NT9. The 5W5V crystal corresponds to TBK1 co-crystallized with a substrate, while 6NT9 corresponds to the TBK1-STING complex. Structural analysis of TBK1 was performed with Molecular Operating Environment (MOE) (33). MOE showcased the amino acids of TBK1 which interacted with ligands or other proteins, such as STING (47). Subsequently, the mutations that were identified in previous steps of the analysis as well as the interaction sites between TBK1 and the ligands were studied carefully. Specifically, each PDB entry was examined for ligand interaction using MOE’s Ligand Interaction module. The results of this analysis, in conjunction with related information extracted from the NCBI Conserved Domain Database, were added onto the consensus sequence in the multiple sequences alignment (48).

## Results

### Dataset

The primary NCBI dataset contained 3.003 entries, related to protein TBK1. This dataset consisted of not only TBK1 protein and its homologs, but also contained unrelated proteins, such as the protein TBKBP1. This irrelevant protein, along with synthetic, hypothetical, partial, low quality and predicted proteins were labeled “noisy” data and were removed from the dataset. Since the NCBI database provides partial duplicate sequences, the sequences with greater than 95% similarity were also removed, retaining the one with the longer length (49). Thus, a final dataset of 283 protein sequences was created. The protein sequences were derived from all kingdoms, from simpler organisms such as Cnidarians, to more complex organisms, such as Homo sapiens. Table 1 shows the phyla from each kingdom, which have been shown to express TBK1 (**Table 1**). Multicellular organisms encoding TBK1 include arthropods (bees), cnidarians (corals), echinoderms (sea urchins) and rotifers. Chordates encoding TBK1 include mammals (homo sapiens), fishes (goldfish), birds (parrots), turtles, amphibians (frogs), lizards, cartilaginous, lampreys, crocodiles, and alligators. TBK1 is not expressed in the kingdoms of bacteria, protista, archaea, alants, fungi and viruses.

**Table 1:**
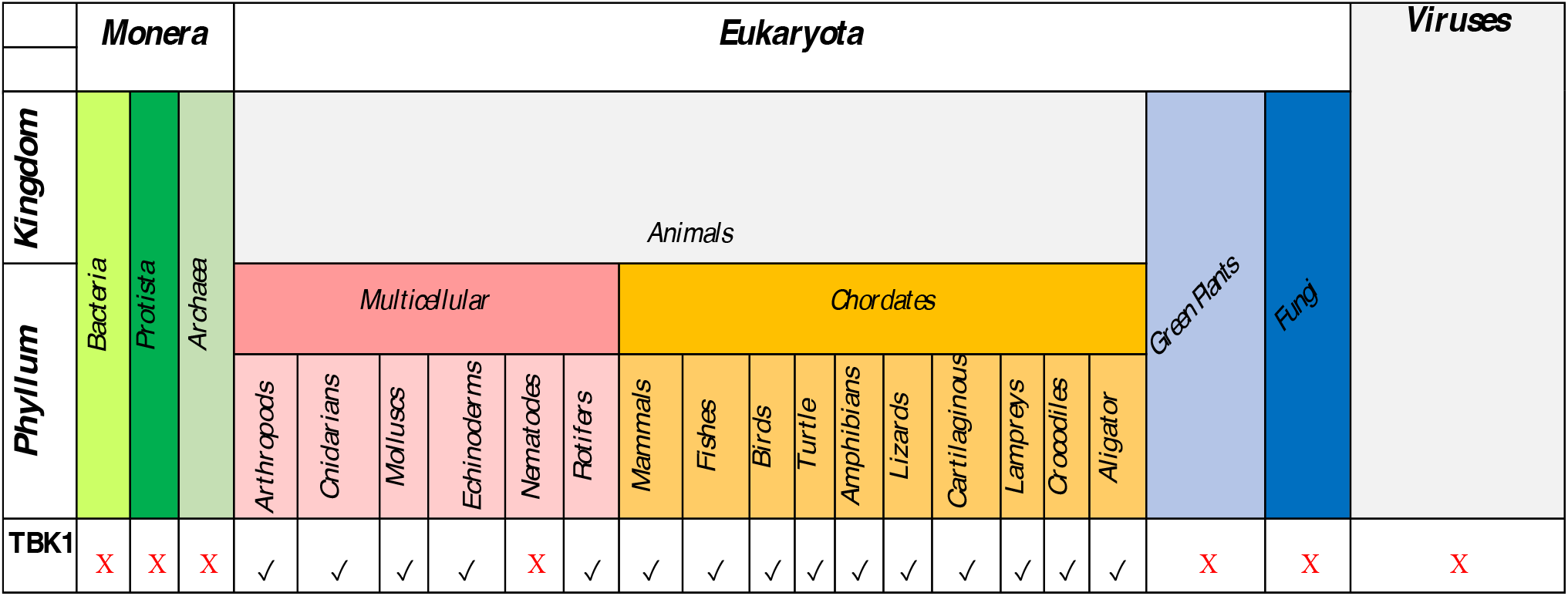
Class groups expressing TBK1, according to which the representative sequences were selected in order to create the final dataset.

### Multiple sequence alignment and conserved motifs

Multiple sequence alignment (MSA) of TBK1 sequences was performed to identify highly conservative regions within all organisms (50). As mentioned, TBK1 harbors conserved functional domains, including kinase KD, ULD and ED, with the latter encompassing CCD1 (LZ and HLH domains) and CCD2 (35). Careful study of the MSA using Jalview resulted in the identification of those domains in most phyla, although a variation of the number of repeats and length is noticeable. In order to compare the human TBK1 with that expressed by other organisms, one species from each class was randomly selected and studied. As previously described, the human TBK1 length is 729, with a 301 aa long KD domain, a 305-383 aa long ULD domain and a 407-729 aa long ED domain. The results are presented in Table 2. The first column lists the classes, the second column shows the organism-representative of each class, and the third column shows the amino acid sequence of TBK1 for each representative organism. The fourth column lists the deviations within these sequences, compared to the human TBK1. For example, echinoderm TBK1 has 57 additional amino acids in its sequence, while reptile TBK1 exhibits 10 amino acids less than human TBK1. In addition, within the table, the number of amino acids that have been removed is shown for each domain. It is interesting to note that there are organisms, such as the turtle, whose TBK1 exhibits no deviation from human TBK1, being exactly 729 aa long, as indicated by a “-” in the fourth column (**Table 2, Figure 1**).

**Table 2:** Comparison of human TBK1 domains against other organisms. One species from each class was randomly selected and studied. The first column lists the classes, the second column shows the organism-representative of each class, and the third column shows the amino acid sequence of TBK1 for each representative organism. The fourth column lists the deviations from human TBK1.

| **Phylum** | **Species** | **Length of TBK1 (aa)** | **Compared to human TBK1** |  |
| --- | --- | --- | --- | --- |
| Echinoderms | Strongylocentrotus purpuratus | 786 | +57 aa | 5 KD<br>52 ED |
| Cnidaria | Acropora millepora | 779 | +50 aa | 1 KD<br>2 ULD<br>47 ED |
| Mollusca | Crassostrea gigas | 769 | +40 aa | 7 KD<br>33ED |
| Rotifers | Brachionus plicatilis | 768 | +39 aa | 8 KD<br>8 ULD<br>23 ED |
| Fishes | Acanthochromis polyacanthus | 744 | +15 aa | 14 KD<br>1 ED |
| Lampreys | Petromyzon marinus | 744 | +15 aa | 15 ED |
| Crocodiles | Crocodylus porosus | 741 | +12 aa | 12 KD |
| Cartilaginous | Amblyraja radiata | 730 | +1 aa | 1 ED |
| Birds | Aquila chrysaetos chrysaetos | 729 | - | - |
| Alligators | Alligator mississippiensis | 729 | - | - |
| Turtles | Chelonia mydas | 729 | - | - |
| Lizards | Lacerta agilis | 729 | - | - |
| Amphibians | Xenopus tropicalis | 725 | - 4 aa | 4 ED |
| Arthropods | Drosophila elegans | 720 | - 9 aa | 9ED |
| Reptiles | Notechis scutatus | 719 | - 10 aa | 10 ED |

One key observation is the large level of conservation exhibited at domains KD and ULD, with small exceptions. In contrast, the elongated helices domain (ED) exhibits strong variation across the different species. Human TBK1 is evolutionarily closer to TBK1 of birds, alligators, crocodiles, lizards, cartilaginous and turtles, and evolutionarily distant to echinoderms, cnidarians, mollusca, arthropods and reptiles.

Aside from the study of evolutionary relationships, the MSA enables the search for motifs. An indepth review of the consensus sequence and TBK1 mutations provides a basis for the identification of six highly conserved motifs (**Figure 2**). A more targeted approach to human TBK1 protein can provide data on its importance for pharmaceutical target. Motif A is marked with green color and encompasses the consensus sequence [HLRENGIVHRDIKPGNIM], residues 125 to 142 (**Figure 2**). Mutations have been marked with blue squares and ligand interactions with red squares. In the 6NT9 crystal of human TBK1, this motif it appears to be next to the active site of protein, close to the complexed ligand. This region exhibits high levels of conservation. Four mutations were identified: R, N, V, R and 1 site of ligand interaction: M. Motif B is marked with red color and encompasses the consensus sequence [YKLTDFGAAREL], residues 153 to 164 (**Figure 2**). In the 6NT9 crystal of human TBK1, this motif is next to the active site of protein. This region exhibits high conservation and only a single mutation was detected: T, which also functions as a ligand interaction. Motif C is marked with brown color and corresponds to the consensus sequence [DDEQFVSLYGTEEYLHPDMYERAVLRK], residues 166 to 192 (**Figure 2**). In the literature, only the DDEQFVSLYGTEE sequence is mentioned as a motif, but in the context of our analysis, it was extended because of its site in the 6NT9 crystal. Specifically, this motif contains the residue Ser172, which is important for the activation of TBK1. This motif exhibits high conservation levels, while two mutations were detected: D, Y, with no detected site of ligand interaction. Motif D is marked with blue color and occupies the consensus sequence [LPFRPFEGPRRNKEVM] residues 219 to 234 (**Figure 2**). In the 6NT9 crystal, this motif appears to be located next to the active site. Two mutations were detected in the otherwise conserved region: R, R, and no point of interaction. Motif E is marked with pink color and occupies the consensus sequence [ENGPIDWSGDMP], residues 253 to 264 (**Figure 2**). This region appears less conserved and three mutations were detected: N, I, P, as well as no point of interaction. On a structural level, this motif is not related to the active site of the protein, however further studies may better elucidate his role. Lastly, motif F is marked with yellow color and corresponds to the consensus sequence [YNEEQIHKFDKQK], encompassing residues 577 to 589 (**Figure 2**). One mutation, Q, was identified, along with seven detected points of interaction: Y, N, Q, I, K, F, Q. In the 6NT9 complex crystal, this motif is adjacent to the STING protein. In conclusion, the motifs A, B, C, D and E were identified within the kinase domain. Only one motif, F, was identified the CCD1 domain, more specifically between the domains LZ and HLH. Given their structural relationship with the active site of human TBK1, these motifs are identified as potential targets for novel therapeutics in the ALS setting.

**Figure 2:**
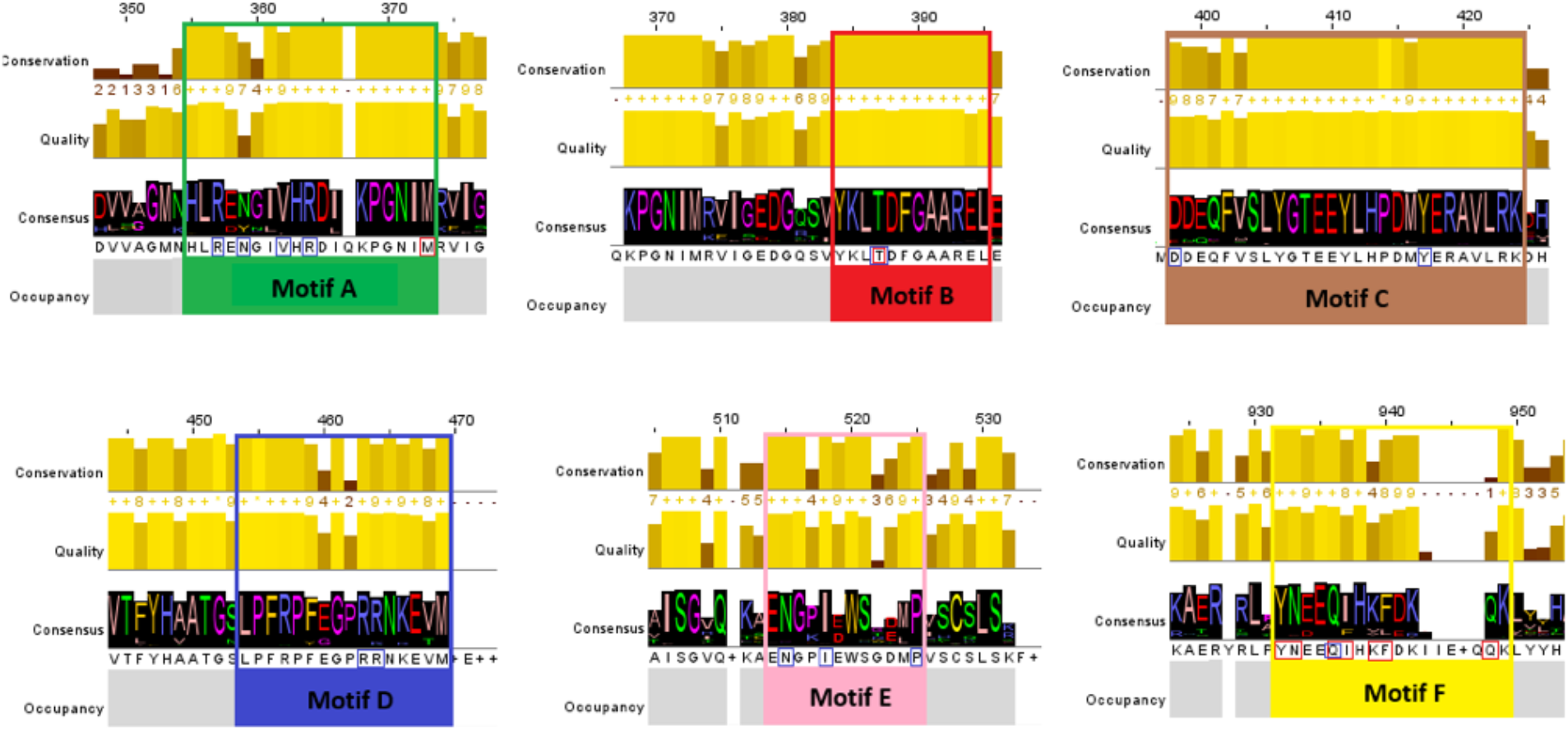
Multiple sequence alignment of TBK1 representative sequences, visualized in Jalview. The consensus sequence of the MSA and TBK1 mutations allow the identification of six highly specific conserved motifs: A, B, C, D, E and F, which may constitute novel pharmacological targets in Amyotrophic Lateral Sclerosis.

### Phylogenetic analysis

Phylogenetic analysis revealed that protein TBK1 was separated into 10 phyletic branches (**Figure 3**). Each branch represents a different phylum of all the species that encode TBK1. The tree appears to be divided into many branches from the root (50). There are 10 phyla represented in the tree: Mammals, Birds, Reptiles, Chordate members, Fishes, Rotifers, Arthropods, Echinoderms, Mollusca, and Cnidarians. The human TBK1 is found in the last branch of the phylogenetic tree and appears to be evolutionarily closer to the TBK1 of birds. However, it seems that the ancestral origin of TBK1 is most likely traced back to the Rotifers (light blue), as they correspond to the first branch that starts from the root of the tree. The following branches, in order, correspond to Mollusca (brown), Echinoderms (red), Arthropods (light green) and the Cnidarians (fuchsia). The phylum of fish follows in light blue. Note that fish include lampreys and cartilaginous for easier identification. Mammals appear at the end of the tree. Various representatives of the wider leaf of the Chordates appear in orange, which seem to be classified in the specific point, because they are partial areas from crystalline structures. The last leaf of Mammals contains humans and organisms evolutionarily closer to humans, such as apes. Reptiles, including crocodiles, alligators, lizards, snakes, and turtles, are observed in the dark green. Finally, the bird branch is marked with pink color (**Figure 3**).

**Figure 3:**
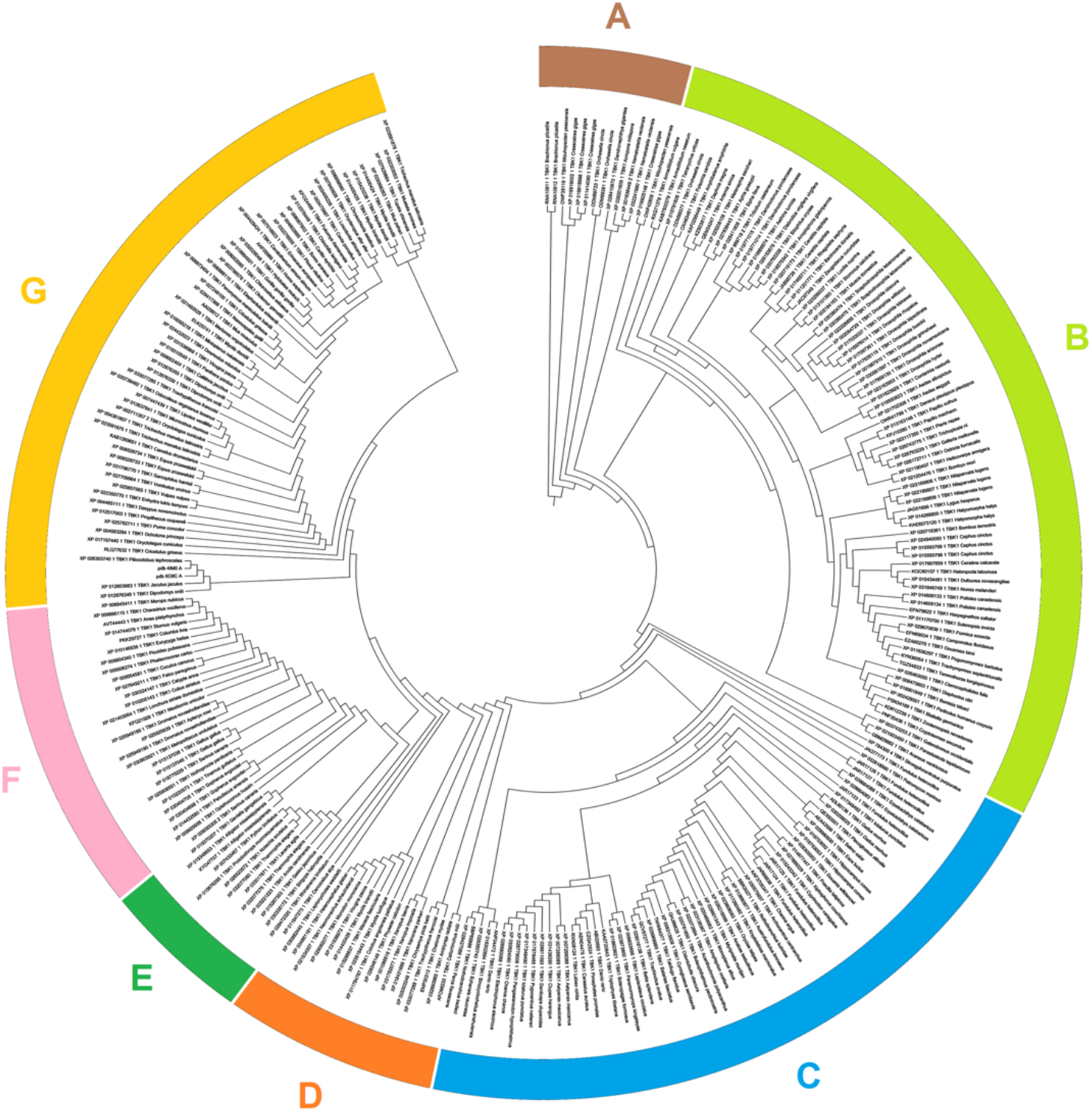
Phylogenetic analysis of protein TBK1, visualized through MEGA software. A: Rotifers, Echinoderms, Cnidaria and Mollusca, B: Arthropods, C: Fishes, D: Amphibians, E: Reptiles, F: Birds, G: Mammals.

### Single nucleotide polymorphisms, variants, and mutation analysis

TBK1 mutations and SNPs were collected from the database GWAS Catalog, and related publications (50). An SNPs and mutation analysis has been performed in order to mapping the TBK1 with the corresponding pathogenic conditions. Afterwards a more specialized analysis has been performed towards to extracting all the available SNPs that are directly connected with the ALS disease (50). Based on the results, they have been identified several types of mutations including (a) Silent mutations (b) Missense mutations (c) Nonsense mutations and (d) frameshift mutations. All the TBK1 mutations are presented in detail in the Table 3. In total, they have been identified 85 mutations in TBK1 protein. The majority of the studied mutations (32 different cases) were detected at the kinase domain (residues 9-301). Moreover, in the ULD domain were detected only 5 mutations, in the linked region (between ULD and CCD1) were detected 7 mutation and in the CCD2 were detected 7 mutations. Finally, at the CCD1 domain (residues 407-657) were detected 27 mutations. Most of the studied cases in mutations were replacements of an amino acid by another amino acid, which eventually leads to a modified TBK1, with a loss of kinase function. In some other cases, silence mutations have a negative effect on the TBK1 protein through several epigenetic factors that are taken place (51). Based on SNPs observations the TBK1protein is involved in several biological pathways including signaling, modification and activation of other proteins, receptors, and cascades, such as TLR4, Notch and IRF3/IRF7 (**Table 4**).

**Table 3:**
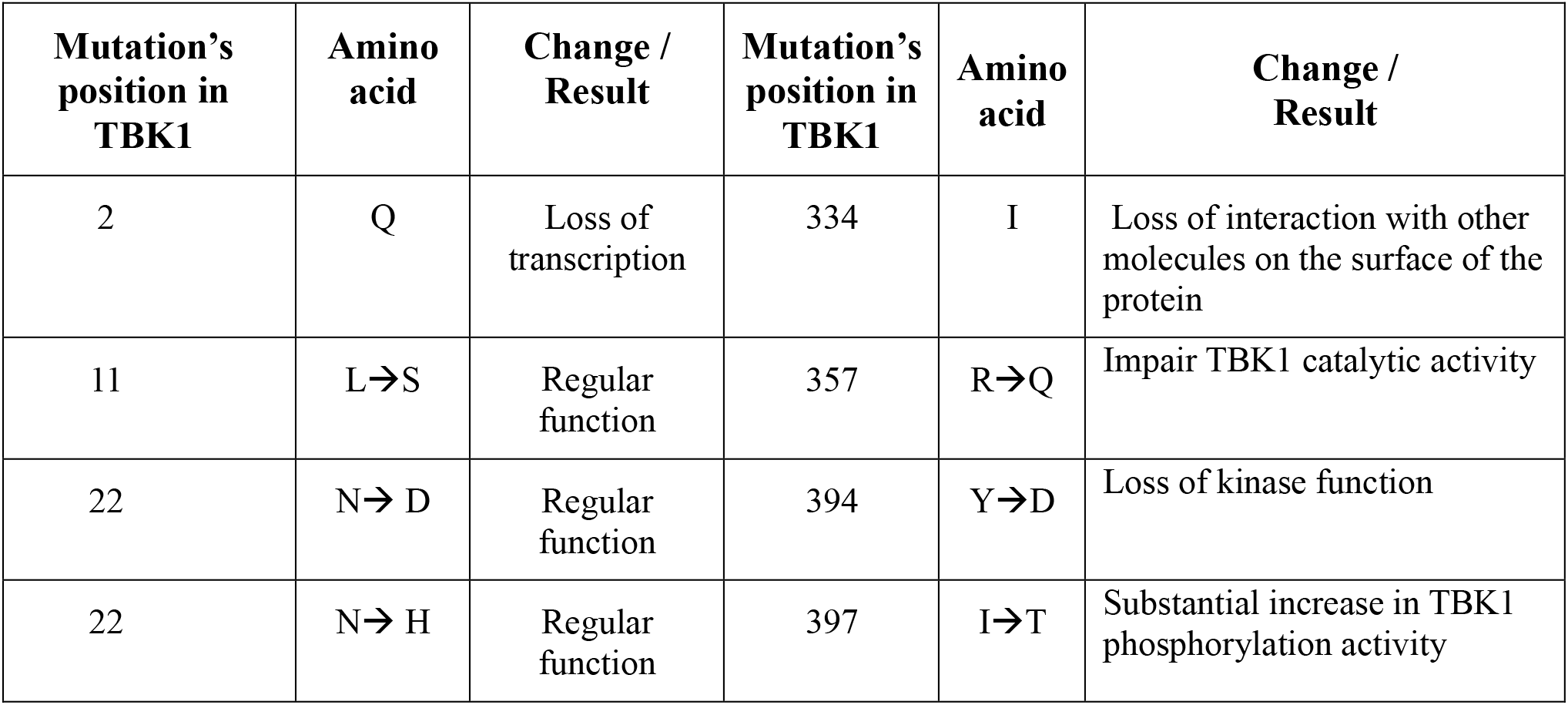
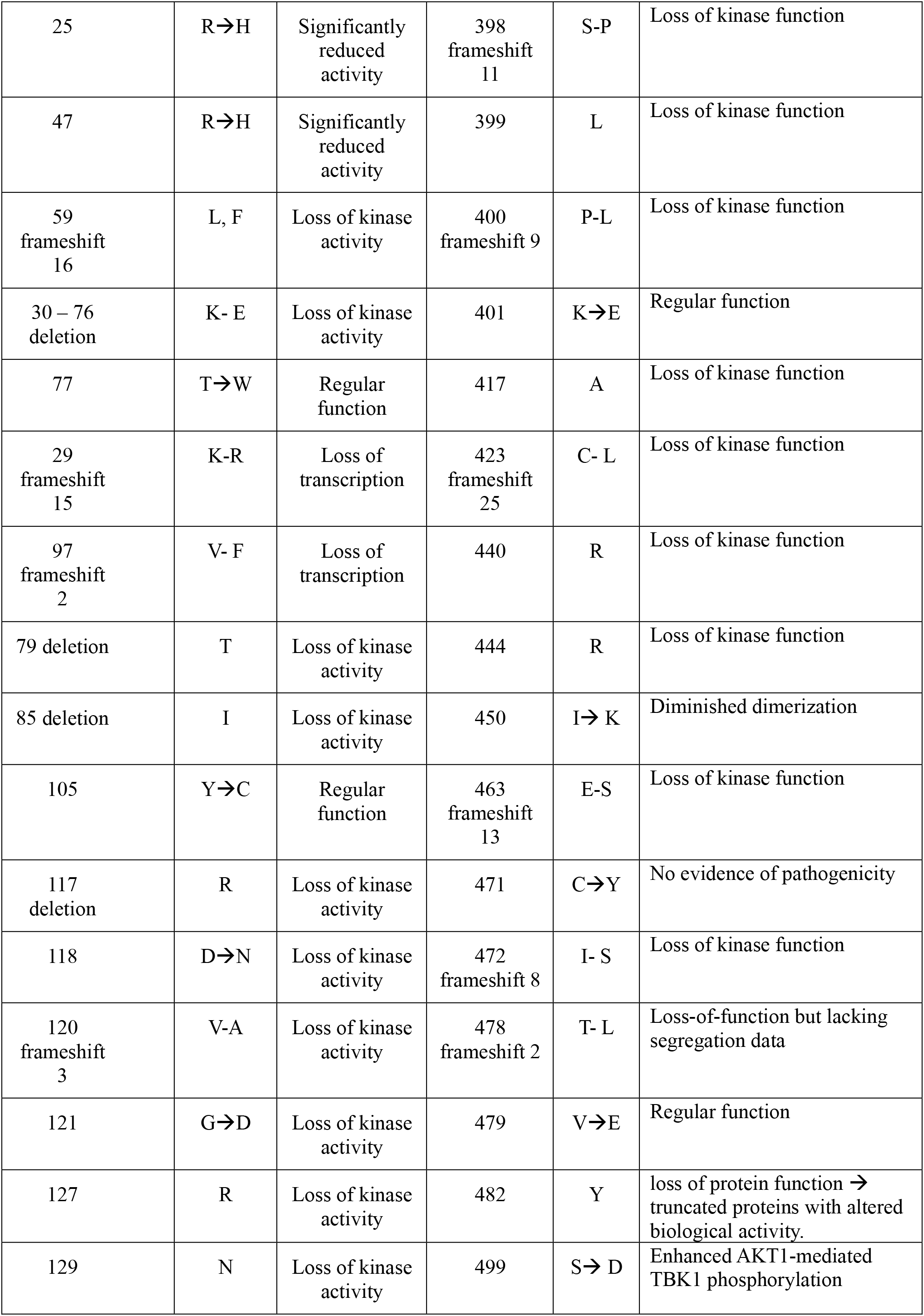
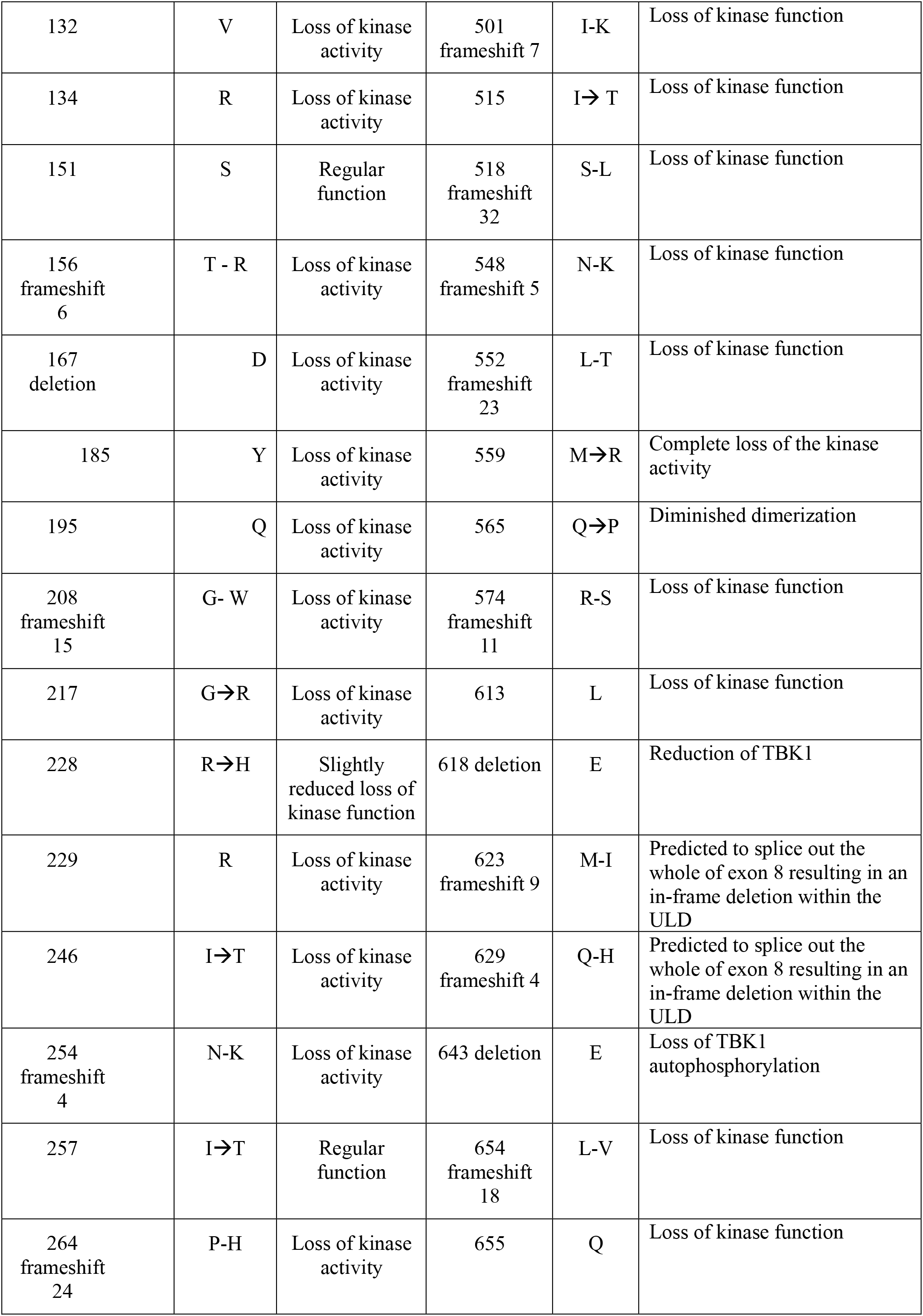
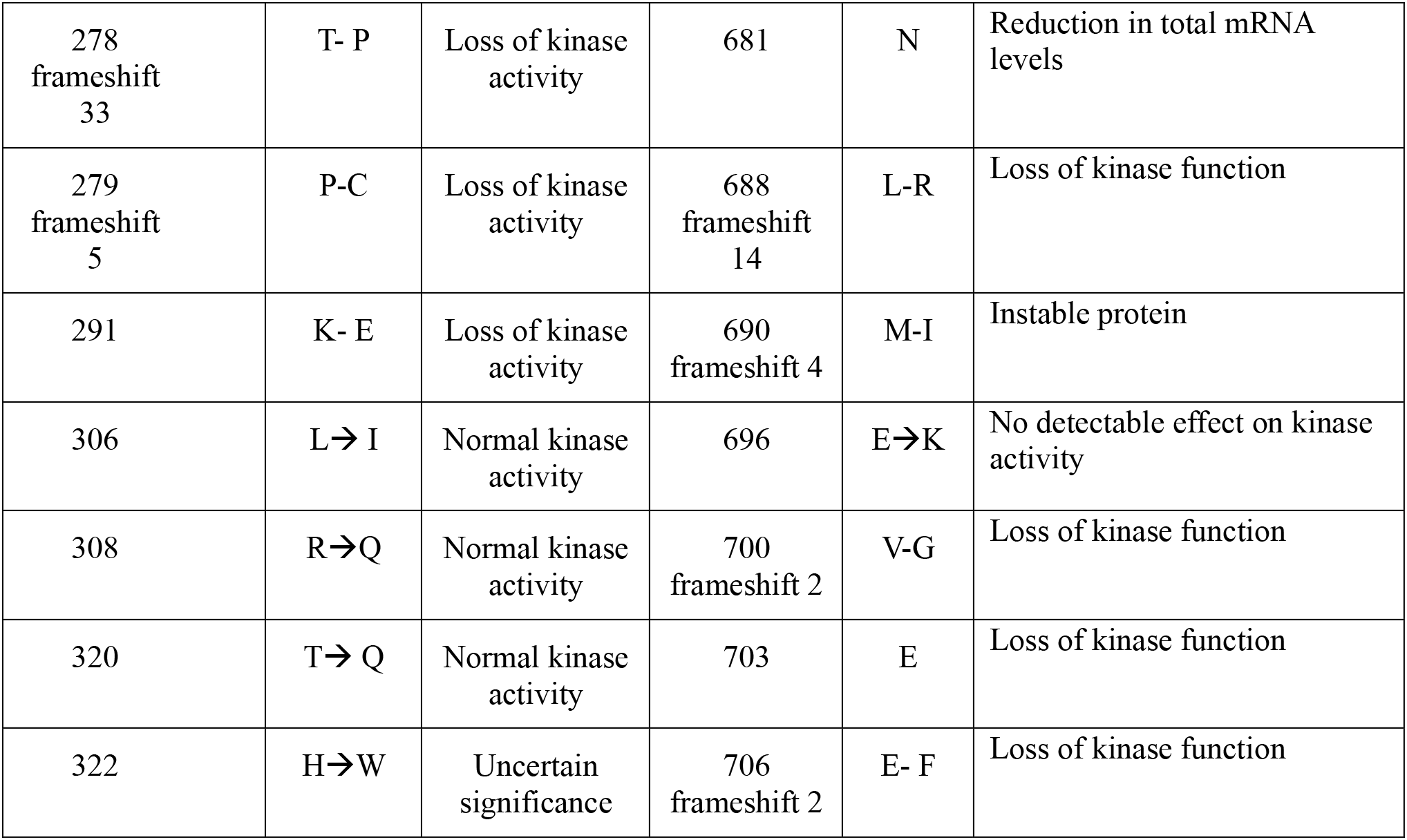
Mutations in TBK1 and their effect.

| Mutation's position in TBK1 | Amino acid | Change / Result | Mutation's position in TBK1 | Amino acid | Change / Result |
| --- | --- | --- | --- | --- | --- |
| 2 | Q | Loss of transcription | 334 | I | Loss of interaction with other molecules on the surface of the protein |
| 11 | L→S | Regular function | 357 | R→Q | Impair TBK1 catalytic activity |
| 22 | N→D | Regular function | 394 | Y→D | Loss of kinase function |
| 22 | N→H | Regular function | 397 | I→T | Substantial increase in TBK1 phosphorylation activity |
| 25 | R→H | Significantly reduced activity | 398 frameshift 11 | S-P | Loss of kinase function |
| 47 | R→H | Significantly reduced activity | 399 | L | Loss of kinase function |
| 59 frameshift 16 | L, F | Loss of kinase activity | 400 frameshift 9 | P-L | Loss of kinase function |
| 30 – 76 deletion | K- E | Loss of kinase activity | 401 | K→E | Regular function |
| 77 | T→W | Regular function | 417 | A | Loss of kinase function |
| 29 frameshift 15 | K-R | Loss of transcription | 423 frameshift 25 | C- L | Loss of kinase function |
| 97 frameshift 2 | V- F | Loss of transcription | 440 | R | Loss of kinase function |
| 79 deletion | T | Loss of kinase activity | 444 | R | Loss of kinase function |
| 85 deletion | I | Loss of kinase activity | 450 | I→ K | Diminished dimerization |
| 105 | Y→C | Regular function | 463 frameshift 13 | E-S | Loss of kinase function |
| 117 deletion | R | Loss of kinase activity | 471 | C→Y | No evidence of pathogenicity |
| 118 | D→N | Loss of kinase activity | 472 frameshift 8 | I- S | Loss of kinase function |
| 120 frameshift 3 | V-A | Loss of kinase activity | 478 frameshift 2 | T- L | Loss-of-function but lacking segregation data |
| 121 | G→D | Loss of kinase activity | 479 | V→E | Regular function |
| 127 | R | Loss of kinase activity | 482 | Y | loss of protein function → truncated proteins with altered biological activity. |
| 129 | N | Loss of kinase activity | 499 | S→ D | Enhanced AKT1-mediated TBK1 phosphorylation |
| 132 | V | Loss of kinase activity | 501 frameshift 7 | I-K | Loss of kinase function |
| 134 | R | Loss of kinase activity | 515 | I→T | Loss of kinase function |
| 151 | S | Regular function | 518 frameshift 32 | S-L | Loss of kinase function |
| 156 frameshift 6 | T - R | Loss of kinase activity | 548 frameshift 5 | N-K | Loss of kinase function |
| 167 deletion | D | Loss of kinase activity | 552 frameshift 23 | L-T | Loss of kinase function |
| 185 | Y | Loss of kinase activity | 559 | M→R | Complete loss of the kinase activity |
| 195 | Q | Loss of kinase activity | 565 | Q→P | Diminished dimerization |
| 208 frameshift 15 | G- W | Loss of kinase activity | 574 frameshift 11 | R-S | Loss of kinase function |
| 217 | G→R | Loss of kinase activity | 613 | L | Loss of kinase function |
| 228 | R→H | Slightly reduced loss of kinase function | 618 deletion | E | Reduction of TBK1 |
| 229 | R | Loss of kinase activity | 623 frameshift 9 | M-I | Predicted to splice out the whole of exon 8 resulting in an in-frame deletion within the ULD |
| 246 | I→T | Loss of kinase activity | 629 frameshift 4 | Q-H | Predicted to splice out the whole of exon 8 resulting in an in-frame deletion within the ULD |
| 254 frameshift 4 | N-K | Loss of kinase activity | 643 deletion | E | Loss of TBK1 autophosphorylation |
| 257 | I→T | Regular function | 654 frameshift 18 | L-V | Loss of kinase function |
| 264 frameshift 24 | P-H | Loss of kinase activity | 655 | Q | Loss of kinase function |
| 278<br>frameshift<br>33 | T- P | Loss of kinase<br>activity | 681 | N | Reduction in total mRNA<br>levels |
| 279<br>frameshift<br>5 | P-C | Loss of kinase<br>activity | 688<br>frameshift<br>14 | L-R | Loss of kinase function |
| 291 | K- E | Loss of kinase<br>activity | 690<br>frameshift 4 | M-I | Instable protein |
| 306 | L→ I | Normal kinase<br>activity | 696 | E→K | No detectable effect on kinase<br>activity |
| 308 | R→Q | Normal kinase<br>activity | 700<br>frameshift 2 | V-G | Loss of kinase function |
| 320 | T→ Q | Normal kinase<br>activity | 703 | E | Loss of kinase function |
| 322 | H→W | Uncertain<br>significance | 706<br>frameshift 2 | E- F | Loss of kinase function |

**Table 4:** Biological pathways in which the *TBK1* gene is implicated.

|  |
| --- |
| L17 Signaling Pathway |
| Notch Signaling Pathway |
| Regulation of toll-like receptor signaling pathway |
| RIG-I-like Receptor Signaling |
| TLR4 Signaling and Tolerance |
| TNF alpha Signaling Pathway |
| Toll-like Receptor Signaling Pathway |
| Activated TLR4 signalling |
| Activation of IRF3/IRF7 mediated by TBK1/IKK epsilon |
| IRF3 mediated activation of type 1 IFN |
| IRF3-mediated induction of type 1 IFN |
| MyD88-independent TLR3/TLR4 cascade |
| Negative regulators of RIG-I/MDA5 signaling |
| Regulation of innate immune responses to cytosolic DNA |
| RIG-I/MDA5 mediated induction of IFN-alpha/beta pathways |
| STAT6-mediated induction of chemokines |
| STING mediated induction of host immune responses |
| Toll Like Receptor 3 (TLR3) Cascade |
| Toll Like Receptor 4 (TLR4) Cascade |
| TRAF3-dependent IRF activation pathway |
| TRAF6 mediated IRF7 activation |
| TRIF-mediated TLR3/TLR4 signaling |
| ZBP1(DAI) mediated induction of type I IFNservous system |

### Structural analysis

The structural analysis of the TBK1 protein has been performed using the available human crystal structures (PDB 5W5V and 6NT9) of the TBK1 and it co-crystalized ligands with the MOE software (52). All the extracted conserved motifs from the previous steps, they have been identified and marked in the studied crystal structures (**Figure 2 and 4**). In addition, the contribution of the conserved motifs to the formation of the active center of the protein was studied, as well as were identified the critical motifs that are necessary for the TBK1 protein in order to interact with other proteins such as the STING protein. The TBK1 active site is formed by fourteen “key” amino acids including Leucine 15, Valine 23, Alanine 36, Valine 68, Methionine 86, Glutamic acid 87, Phenylalanine 88, Cysteine 89, Proline 90, Cysteine 91, Glycine 92, Threonine 96, Methionine 142 and Threonine 156. Based on the available crystal structures, the TBK1 protein is taken place a dimer form, from which each monomer is found to interact with the STING protein (PDB: 6NT9) (47).

**Figure 4:**
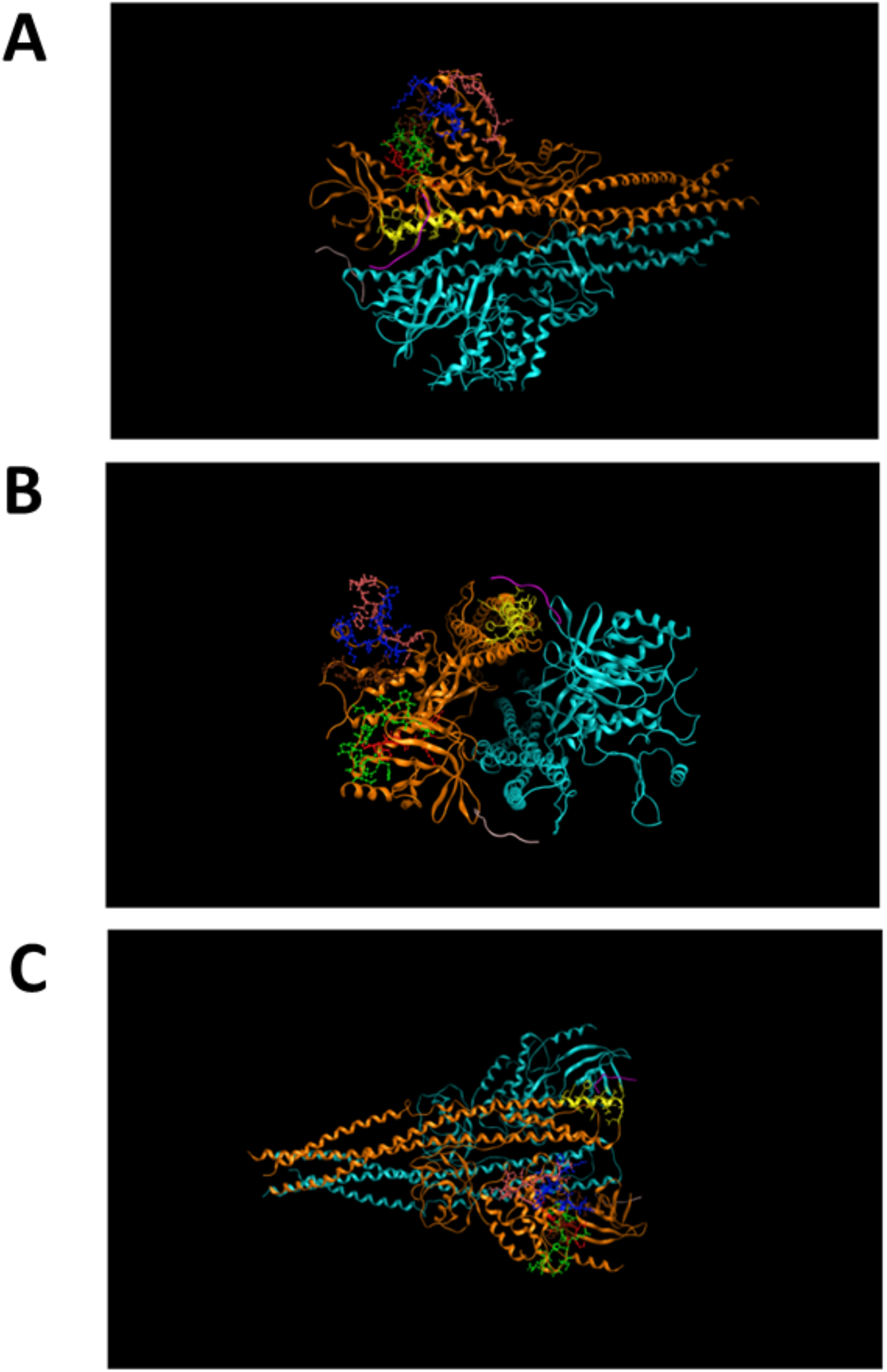
The highly conserved motifs A, B, C, D, E and F mapped onto the TBK1 crystal (PDB ID: 6NT9) in ribbon conformation using MOE. The first TBK1 monomer is shown in blue and the other TBK1 monomer in orange. The STING protein, which is in the active center, appears purple. In green is the highly conserved motif A. In red is the highly conserved motif B. In brown is the highly conserved motif C. In dark blue is the highly conserved motif D. In pink is the highly conserved motif E. In yellow is the highly conserved motif F. **A:** View 1, **B:** View 2, **C:** View 3.

Several conserved motifs that are identified in the present study, are necessary for the interaction of the TBK1 with the corresponding substrates. As an example, mutations in the conserved motif A in the amino acids R127, N129, V132 and R134 can be causes the loss of TBK1 transcription, the loss of TBK1 autophosphorylation and the loss of IRF3 phosphorylation. In motif B was detected one mutation (T156) which leads to loss of kinase function in TBK. In motif C were identified two mutations (D156 and Y185), which according to the literature those mutations lead to the loss of TBK1 auto-phosphorylation. In motif D were detected 2 mutations (R228 and R229) which have been related with the slightly reduced loss of kinase function. Three mutations have been identified in the Motif E including N254, I257 and P264, which lead to the loss of kinase function. Last but not least, in the Motif F was identified one mutation Q561 which leads to the loss of TBK1 auto-phosphorylation. Moreover, Motif F is necessary in order to TBK1 protein interact with the STING protein. In conclusion, the motifs A, B, C, D and E are in the kinase domain-KD. Only the F motif identified in the CCD1 domain and specifically between the domains LZ and HLH (30–32, 53–56) (**Figure 5**). Around the active site, have been identified the highly conserved motifs A, B, C, D, E, and F, hoping that they will serve as novel pharmacological targets, where a future drug will be linked for the cure of Amyotrophic Lateral Sclerosis (40, 48) (**Figure 4 and 5**).

**Figure 5:**
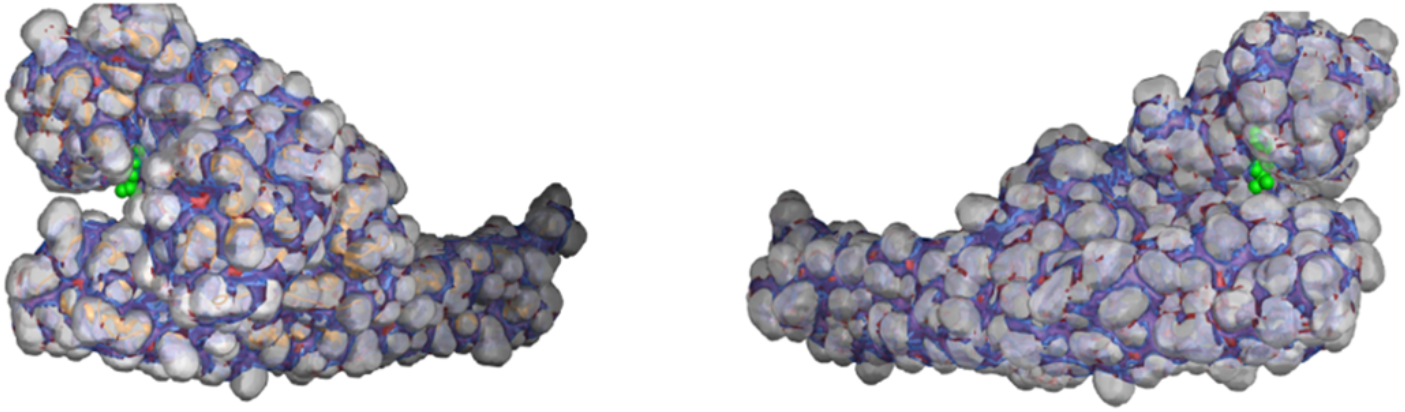
Crystal of the human TBK1 (PDB ID: 5W5V) in MOE. Amino acids set to appear as bubbles, the active site of the protein is in the recess and the ligand appears in green. The highly conserved motifs A, B, C, D, E, and F have been mapped around the active site. These motifs and their structural positions can inform novel therapeutic strategies against Amyotrophic Lateral Sclerosis.

The role of the protein TBK1 as kinase enzyme is to is to interact with several substrates. Kinase proteins can regulate enzymatic and cellular function and for this reason they transfer phosphates on other proteins. Specifically, protein TBK1 belongs to kinases that add phosphate to threonine and serine residues. These amino acids residues have the specific group OH, which can be phosphorylated. Many cellular functions are activated through the protein’s phosphorylation, including as apoptosis, gene expression, membrane transport and proliferation (57). During the reaction of the protein, the active site can reduce the energy of activation, while it increases the rate of reaction. A ligand can bind at the binding site of the protein. The binding site can be included into the active site. Therefore, the binding site catalyzes the binding of a ligand to a molecule and the active site catalyzes the protein’s chemical reaction (58)

## Discussion

The protein TBK1 is important in many biological pathways. It is involved in signaling pathways of other proteins including L17, Notch, TLR4 and TNFa, in the activation of IRF3 and IRF7, in the induction of chemokines, and in the STING protein play a critical role in Toll – like receptors cascades. On the other hand, many mutations have been identified in protein TBK1 that they are leading to important human neurodegenerative disorders, such as ALS. In most of the cases, mutations in TBK1 leads to the loss of kinase activity and diminished dimerization. In the cases where the TBK1 protein is deactivated or damaged, then the signaling pathways are impossible to take place and this leads to the occurrence of several diseases in an organism.

In this present study, a holistic evolutionary analysis of TBK1 protein was performed and conserved motifs that may play a critical role in the TBK1 interactome were identified. Moreover, the outputs have been examined under the aspect of the well-known SNPs and mutations, and structural information based on the available protein 3D structures. The ultimate goal of the present study was to combine all the knowledge produced from protein sequences analysis, mutations and structures analysis towards to extracting beneficial knowledge about TBK1 “key” regions in an effort to fight incurable diseases such as Amyotrophic Lateral Sclerosis.

## Acknowledgments

Not applicable.

